# Pose analysis in free-swimming adult zebrafish, *Danio rerio*: “fishy” origins of movement design

**DOI:** 10.1101/2023.12.31.573780

**Authors:** Jagmeet S. Kanwal, Bhavjeet Sanghera, Riya Dabbi, Eric Glasgow

## Abstract

Movement requires maneuvers that generate thrust to either make turns or move the body forward in physical space. The computational space for perpetually controlling the relative position of every point on the body surface can be vast. We hypothesize the evolution of efficient design for movement that minimizes active (neural) control by leveraging the passive (reactive) forces between the body and the surrounding medium at play. To test our hypothesis, we investigate the presence of stereotypical postures during free-swimming in adult zebrafish, *Danio rerio*. We perform markerless tracking using DeepLabCut, a deep learning pose estimation toolkit, to track geometric relationships between body parts. To identify putative clusters of postural configurations obtained from twelve freely behaving zebrafish, we use unsupervised multivariate time-series analysis (B-SOiD machine learning software). When applied to single individuals, this method reveals a best-fit for 36 to 50 clusters in contrast 86 clusters for data pooled from all 12 animals. The centroids of each cluster obtained over 14,000 sequential frames recorded for a single fish represent an *apriori* classification into relatively stable “target body postures” and inter-pose “transitional postures” that lead to and away from a target pose. We use multidimensional scaling of mean parameter values for each cluster to map cluster-centroids within two dimensions of postural space. From a *post-priori* visual analysis, we condense neighboring postural variants into 15 superclusters or core body configurations. We develop a nomenclature specifying the anteroposterior level/s (upper, mid and lower) and degree of bending. Our results suggest that constraining bends to mainly three levels in adult zebrafish preempts the neck, fore- and hindlimb design for maneuverability in land vertebrates.

## Introduction

Behavior can be both complex and/or consist of relatively simple and stereotypic movement patterns [Saint-Amant, and Drapeau, 1998; Korn, and Faber, 2005]. These movement patterns can be dissected into a sequence of body trajectories in egocentric and allocentric coordinates that can extend over multiple timescales. Examples of relatively simple, stereotypic movements include breathing, which has a major autonomic component, and chewing. Locomotion is another common but more complex behavior of bodily movement, involving alternating, rhythmic contractions of wing muscles for flying in aerial species, and limbs for walking in land vertebrates. The behavioral space in time and across the three spatial dimensions can be vast. For example, primates can express multiple trajectories with limited degrees of freedom at just three joints in their forelimbs to reach the same target [Prablanc et al., 2003; Scott, 2012]. In aquatic species swimming consists of alternating, rhythmic contractions of the axial musculature, and of fins for stabilization. In aquatic species, action patterns are relatively low dimensional and generally restricted to turning and swimming movements. In teleosts, these are largely accomplished by localized and sequential contractions, respectively, of a series of W-shaped muscle bands along the flank [Mathis et al., 2018]. A different magnitude and rate of rhythmic contractions of these muscles as well as undulations of the tail-fin contribute to swimming at different speeds [Severi et al., 2014; Plaut, 2000; Rival et al., 2021].

Zebrafish (*Danio rerio*) were selected as a vertebrate model organism because they are small, hardy fish that lay large numbers of externally fertilized eggs, and the rapidly developing transparent embryos facilitate the identification of induced mutant phenotypes for genetic screens. An exquisite analysis of embryonic development and a large collection of molecular tools, including sequencing of the zebrafish genome, have contributed to the success of this model. As vertebrate organisms showing an ∼70% similarity in their genetic make-up to humans, zebrafish inform most aspects of human biology including the presence of neurological and movement disorders, such as Parkinson’s disease [Howe et al., 2013]. Although behavioral and neural complexity increases with age, neuroscience studies have predominately focused on larval zebrafish, partly because their small size and transparent skull allows visualization of brain activity using two-photon imaging of Ca^2+^ dynamics resulting from neural activity in transgenic larvae. Fine details of movement patterns have also been quite extensively studied in zebrafish larvae [Sumbre, and de Polavieja, 2014; Roussel et al., 2021].

These studies show that seven days post hatching, larvae swim in a ‘‘burst and glide’’ fashion, propelling themselves by short bursts of tail movement, or swim bouts, alternating with interbout periods, where the fish only moves passively through the water. The bouts are themselves constructed from a basic set of tail movements and are executed in sequences specific to different behaviors [Marques et al., 2018]. Studies classifying postures, maneuvers and swim patterns in adult zebrafish, however, are lacking, although a lot of work has been done on the undulatory waves of contraction and relaxation of the muscle segments or myomeres for swimming. One reason for this is that swim patterns in adults are complex and difficult to dissect and describe on the basis of human observation alone.

Swimming and other natural body actions can be thought of as a sequence of continuous and complex sequences of postures articulated by motion [Jellema, and Perrett, 2005] and may be decoded as such both by the individual initiating the action, and by an observer without witnessing the entire action sequence. [Jellema, and Perrett, 2005; Perrett, 1999]. Since articulated action involves initiation of muscle contractions at select musculoskeletal loci leading to targeted vs. transitional postures in space-time, a study of postures and postural transitions can deepen our understanding of the biomechanical constraints for movement design, including its neural control. In principle, the spatial and computational complexity of a continuum of trajectory spaces is vast. One strategy to simplify this is to break it down into a discrete set of elements within postural space where sequences of postures are linked by passive motion governed by biomechanical and hydrodynamic constraints within transitional space. Zebrafish larvae exhibit only 3 basic postures during free-swimming activity [Girdhar et al., 2015]. Their swimming patterns can be of fifteen types, thirteen of which were observed (struggles and S-starts were not observed) using unsupervised clustering methods [Marques et al., 2018]. These swimming patterns are used flexibly in various combinations across different behavioral contexts and their classification can be used to dissect the sensorimotor transformations underlying larval social behavior and hunting. There are no data yet on postural configurations and their classification in adult zebrafish.

Here, we hypothesized that free swimming activity in adult zebrafish can be expressed as sequence of a relatively small number of target poses. In this scenario, continuous motion can be thought of as smooth transitions in space-time between a small set of target poses in contrast to a smooth wave of increasing amplitude traveling from the head-to-tail [Lauder et al., 2011]. Pose transitions are a natural outcome of the physics of bodily mechanics, stiffness and its interaction with the surrounding aquatic medium. Transition phases are unlikely to be actively (neuronally) controlled and maintained. Others have regarded these transitions as unobservable Markov chain processes or states [Li et al., 2013]. To test our hypothesis, we tracked free-swimming movements in adult zebrafish and used a machine-learning, pose-estimation toolkit, DeepLabCut [Mathis et al., 2018] followed by unsupervised clustering for classifying target poses as static expressions of an end-of-trajectory pause. A momentary pause can result either from a preprogrammed phase during the initiation of a trajectory, or be determined by proprioceptive stop-action feedback, or represent the end of a “fixed action pattern” [Ronacher, 2019].

Here, we identify, describe and classify stereotypical postures and their transitions in free-swimming adult zebrafish. This requires precise measurement and a quantitative description of movement co-ordinates of multiple body parts as facilitated by recent applications of machine learning methodologies [Mathis et al., 2018; Mirat et al., 2013; Franco-Restrepo et al., 2019; Romero-Ferrero et al., 2019]. We report here the identification of a relatively small number of target postures that zebrafish exhibit when swimming under semi-natural conditions. We suggest that despite a seemingly uniform organization of flank musculature, movement control in fish is largely localized to or dominated by muscle contractions occurring at specific antero-posterior levels. This can be translated to a “neck, forelimb and hindlimb-centered” design of movement control present in land vertebrates. Our results provide new insights from the viewpoint of postural design of a bauplan for the organization of locomotion and its control in land vertebrates.

## Materials and Methods

### Animal acquisition and maintenance

Wildtype zebrafish (*Danio rerio*) were bred and reared in a 14:10 light:dark cycle at 28°C within the institutional zebrafish facility. At early larval stages, they were fed daily with brine shrimp and as juveniles/adults with specially prepared dry flake food. Each 2.5-liter tank filled with filtered and aerated flowing habitat water typically contained 5 males and 5 females. We raised a cohort of ∼50 wildtype zebrafish. For experiments, fish were transported to the laboratory in a small tank containing habitat water. Care was taken to keep fish in the same habitat water during all transport and handling procedures. For behavioral observations, individual fish were gently scooped out of the holding tank with a small container together with habitat water to avoid out-of-water exposure and any stress resulting from transient hypoxia and handling. Twelve 3-to-10 months old (7 males, 5 females) juvenile/adult zebrafish were selected at random for this study. All procedures were approved by the Institutional Animal Care and Use Committee.

### Recording apparatus

Adult zebrafish were gently introduced, within a small (17 x 17 cm) tank under relatively uniform room lighting. The tank had a smooth, curved, and white, flat bottom and sides. A webcam (Logitech C922) was placed ∼30 cm above the tank to ensure that the entire tank was always within its field of view. Water depth was adjusted to nearly 2 cm to minimize distress, while keeping the fish within the depth of focus of the camera. The data obtained therefore corresponds to near-surface swimming behavior within a two-dimensional plane in adult zebrafish. Video recordings were initiated immediately after fish placement and continued for 10 minutes or less at 30 frames/second (fps) using LogiCapture video capture software. For some videos, an Okiocam document camera and software was used. Video files were stored on a desktop computer’s hard drive for off-line analyses.

### Markerless tracking

The body-axis can be defined by two fixed points on the body, such as the anterior-most “’snout” region or the nose of the fish and the “head” region at the center of the skull. The heading direction is determined by the orientation and trajectory of the snout-skull axis relative to the reference frame of the tank. Whereas the snout and head are connected by a relatively inflexible bony skull, a third location on the body axis, the “tail” region at the base of the caudal fin, provided useful information on the angular orientation of the body relative to the snout-skull axis. Different muscle groups determine the relative position and trajectories of important locomotory structures, such as the flank and fins of a zebrafish. To track movement, we focused on those body parts that were easily visible for tracking purposes and that played a key role in initiating characteristic postures. To identify stereotypic static postures, such as body bends required for turning as well as other behavior elements and postural patterns associated with swimming, we also continuously tracked the x-y coordinates of the flank’s midpoint, the base of the caudal fin, and the midpoint of the caudal fin. The base of the tail fin correlates with turning and undulating movements of the tip of the tail fin propels the fish forward.

Videos of single free-swimming zebrafish were analyzed using the machine learning (ML) markerless pose estimation toolkit DeepLabCut (DLC) [Kane et al., 2020; Mathis et al., 2018]. A DLC model was trained with 290 pre-labeled frames of free-swimming zebrafish locomotion for 800,000 iterations on an NVIDIA GeForce GTX 1070 GPU to yield reliable tracking across all frames of a recorded video. Tracking data were normalized across all twelve animals to account for variations in tank displacement and fish size across samples. DLC outputs a series of plots for the locations of the markers throughout the video as well as likelihood plots of accuracy across all frames. It provides a spreadsheet of co-ordinate values of all markers and model accuracy. Graphs of the zebrafish’s plotted coordinates showcased movement patterns across different regions of the tank as the fish engaged in an exploration of its surroundings.

We utilized DLC to track the midpoint between the eyes (ocular center), skull base, upper (level of pectoral fin), mid and lower (base of dorsal fin), the tail-base (base of the caudal fin), tail-mid and tail-tip (Fig. 1 inset). In our initial tracking experiments, we used 10 virtual markers for tracking movement of different body parts. The dorsal and pectoral fins play an important role in stabilizing the body poses, and the base of the tail-fin correlates with turning. Moreover, these tended to frequently overlap with the body, affecting the accuracy of their tracking. Therefore, we avoided marking and tracking the movements of the dorsal and pectoral fins for this study on postural configurations of the body. The middle of the tail-fin region propels the fish forward and both, the sinusoidal undulations of the tail-fin -mid and -tip do not contribute in any significant way to any unique body posture. Also, the tip of the tail-fin was not always easily visible under water owing to its relative transparency and tracking coordinates for this were not used in the analysis.

**Figure 1.**
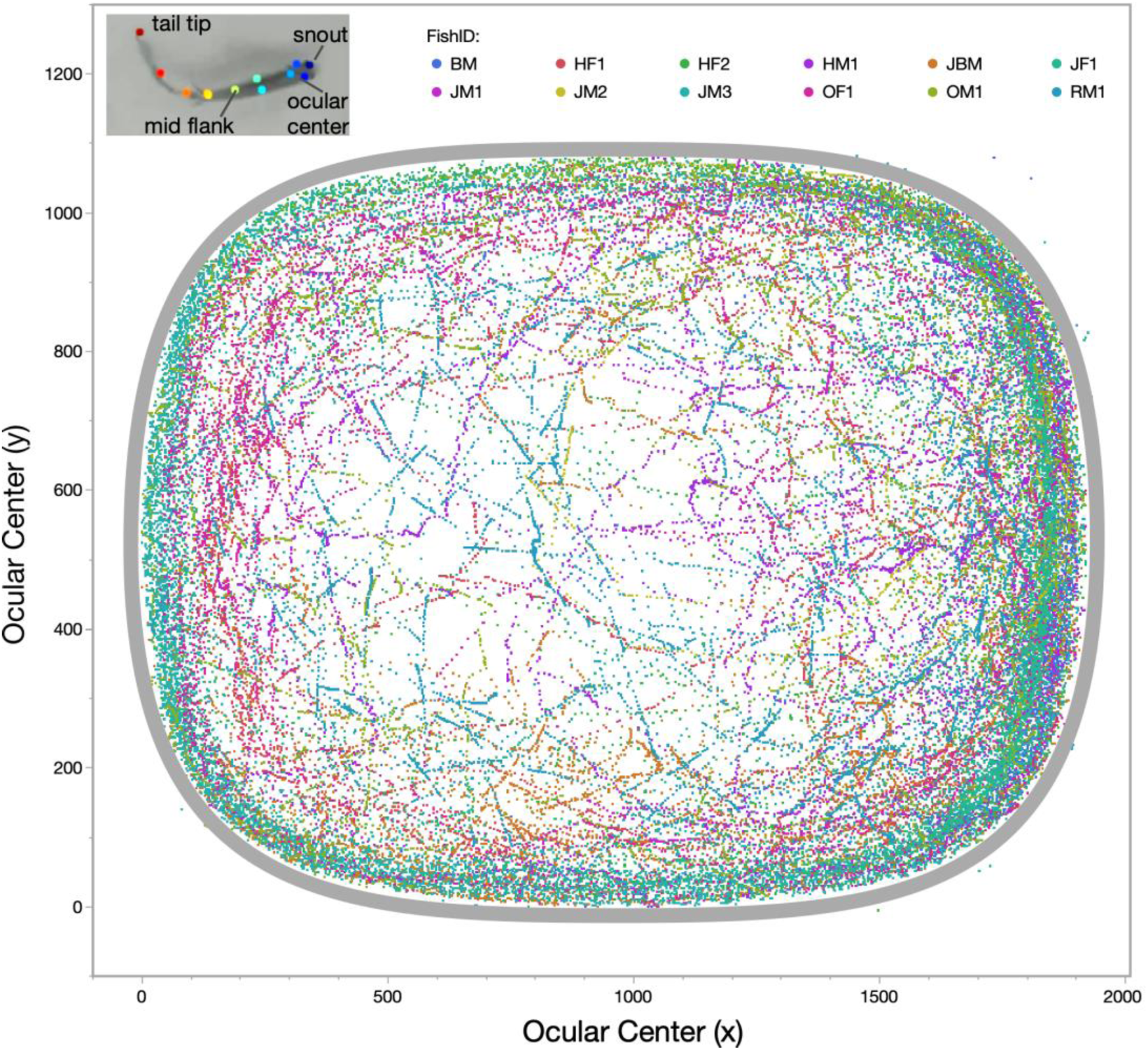
Scatterplot of x-y coordinates showing the swim pattern of 12 adult zebrafish tracked over 5 minutes in the tank arena (grey boundary) using a virtual marker placed at the center of the head between the two eyes (ocular center). A few points outside the boundary of the tank represent frames where tracking failed. Inset shows 11 marker locations on the body of an adult zebrafish that were originally tracked.

### Clustering methods

Discrete, repeated patterns were extracted from tracking data through unsupervised classification with the recently developed B-SOiD software (vers. 2.0)[Hsu, and Yttri, 2019]. B-SOiD pulls repeated behaviors from tracking data generated by DeepLabCut. The B-SOiD algorithm calculates each body part’s displacement, relative distances, and relative angles before performing dimensionality reduction with UMAP. Units of behavior are extracted through unsupervised clustering with HDBSCAN and presented as a time series.

A B-SOiD model was created using nine characteristic videos of individual free swimming zebrafish behavior. For a detailed analysis of static postures in a single adult zebrafish, we marked and continuously tracked the x-y coordinates of eight body parts (including level of snout or ocular center, pectoral fin, pelvic fin, dorsal fin, mid-flank, tailfin-base, tailfin-mid and tailfin tip; see inset in figure 1) for pose-estimation and behavior analysis. For this study, our interest was in B-SOiD-identified “bouts” obtained for frame sequences in the video of free-swimming adult zebrafish. Bouts with a bout-length of 1 frame were selected. For each single-frame motion capture, we calculated multiple parameters, such as distances and angles between markers that were output as a spreadsheet and imported into Microsoft Excel. To overcome inconsistencies in pose classification, revealed via the t-distributed stochastic neighbor embedding (t-SNE) method, we performed a normal mixtures clustering using principal clusters analysis (PCA) of relevant features, such as angles and distances between markers.

### Statistical measures and analyses

For a statistical analysis of our pre-processed data, we used JMP-Pro (vers. 17; SAS Inc.) software. We performed factor analysis to estimate the total variation that could be captured via combinations of features obtained from body position markers. We used principal components analysis to estimate the best parameter combinations to employ for clustering and multivariate analyses, including multidimensional scaling (MDS) to visualize their proximity patterns in 2D-space.

## Results

### Postural clusters pooled across multiple animals

Each individual exhibited free swimming movements across most of the tank area that involved frequent turns, requiring bending of the vertebral column (Figure 1). The B-SOiD toolkit extracted features from x and y coordinates from body flexions during swimming and provided clusters of similar action patterns across multiple time segments via implementation of machine learning approach to perform a multivariate time series analysis. For each frame, the B-SOiD algorithm calculated 6 distances, 6 angles, and 6 displacements between body parts. The output files from B-SOiD analysis contained time-series data with the behavior pattern corresponding to each frame, bout length data, and transition matrices. After testing several B-SOiD parameters (Supplementary Table 1; from B-SOiD Run Summary-update), a model was generated with a minimum cluster size of 0.5% to yield 34 clusters. This model indicated that just four points along with features (body part angles and distances between pairs of points) were enough to capture most of the variation in the geometric relationships between body parts. We applied the model to extract behavior patterns from all 12 fish video recordings.

For postural analysis, we focused on the shortest B-SOiD-classified “bouts” captured within a single frame representing 33.3 ms. B-SOiD reported instances of 34 patterns or pattern IDs for single frames. A visual analysis of frames with same pattern ID, however, indicated that the frames included in each pattern ID captured disparate body postures accounting for a large amount of variation. To test for fidelity of pattern IDs, we conducted discriminant function analysis (DFA) with “pattern ID” as the categorical variable to obtain a quantifiable estimate of the variation captured and overlap across different groups (Figure 2A). Although the first two canonical dimensions of the DFA conducted for pattern IDs captured 90% of the pose variation in individual animals and showed significant separation (P < 0.0001), it revealed a misclassification of nearly 90% of pattern IDs (quadratic fit). Across all animals, the misclassification became worse (> 95.16%). Areas under the ROC curve for pattern IDs ranged from 0.59 to 0.77. A t–SNE plot using normal mixtures clustering algorithm with data points labeled for pattern ID (Supplementary Figure 1). A section of t-SNE plots obtained for shorter time window of 5 minutes revealed more clearly a fractionation of pattern IDs across different clusters of poses (Figure 2B). These results made us question the validity of B-SOiD as a multivariate time series classifier of behavioral patterns despite its reputation as a powerful algorithm among the recently available packages using ML methods. One possible reason may be that the software does not perform well when the classification is restricted to single frames for pose analysis. Another possibility is that the unidimensional body-axis of a fish does not provide a rich multidimensional framework compared to species exhibiting multidirectional limb trajectories for B-SOiD to work well.

**Figure 2.**
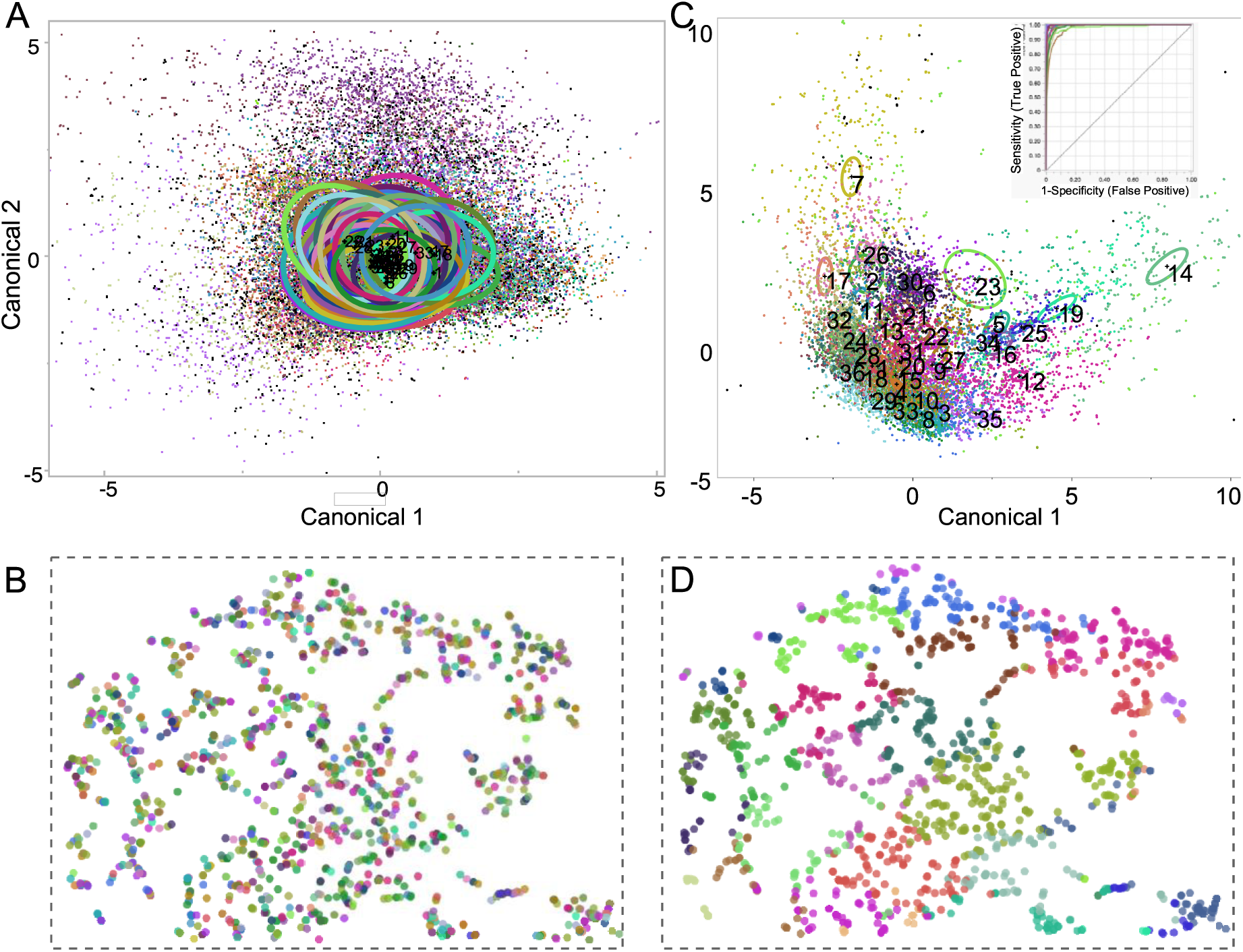
A. Canonical plot for DFA of 34 clusters. B-SOiD classification of single frames taken from a video of the data pooled from all (n = 12) free-swimming adult zebrafish Ellipses for each cluster are for 50% confidence interval boundaries. B. A portion of the 2D t-SNE plot with the B-SOiD-based pattern IDs used for color coding each frame. Placement of the curves away from the diagonal indicates a higher reliability of fit. C. DFA of the feature space for one individual (“BM”) with PCA-generated cluster ID as the color code. Ellipses indicate cluster centroids means of confidence limits. DFA showed 20% misclassification and areas under the ROC curve ranged from 0.9782 to 0.9999 (see inset). D. The same section (as in B) of the 2D t-SNE plot but with normal mixtures clustering IDs used for color coding each frame. Note the proximity of similar colored dots in D compared to those in B, indicating a more consistent clustering of frames with similar poses.

To improve our estimate of postural configurations from that provided for single frames by B-SOiD, we first removed outliers in tracked x-y coordinates from further analysis using a robust fit screening algorithm. We used Excel to compute distances between markers and angles between body segments from x-y coordinates. We once again conducted DFA of new clusters obtained via a normal mixtures clustering algorithm (JMP-Pro, vers. 17; SAS Inc.) with cluster ID as the categorical variable (Figure 2C). An ROC plot for each pattern ID obtained with DFA revealed a low hit and miss rate (Inset in Figure 2C). The DFA was based on 6 parameters identified as the most uncorrelated via factor analysis. These included three distances, namely, dist(OC-MF), dist(OC-TB), dist(MF-TB), and three body angles, angle(OC-MF-TB), angle(MF-TB-TM), angle(OC-MF-TM). Distances between markers contributed maximally to the variation. A t-SNE plots of the same section as in figure 2B revealed a more contiguous distribution of cluster IDs across different clusters of geometric configurations (Figure 2D).

Results from PCA sampled from 3 to 600 clusters revealed a best classification into 86 clusters for x-y coordinate feature space across all 12 zebrafish whose behavior was tracked with DLC over a roughly 10-minute period (Figure 3). We also estimated cluster fits individually for each of the twelve fish. The clustering provided a best fit for a somewhat different range of clusters for each animal, but were typically in the range of 37 to 70 clusters. Although the best fit for all individuals pooled together showed a slightly better fit for 200 and even 600 clusters, the majority of these were virtual clusters with none to < 0.007 % of frames or poses included in each cluster. Only 38 of the 400 to 600 clusters included a population that included a => 0.01% proportion of the total number of frames.

**Figure 3.**
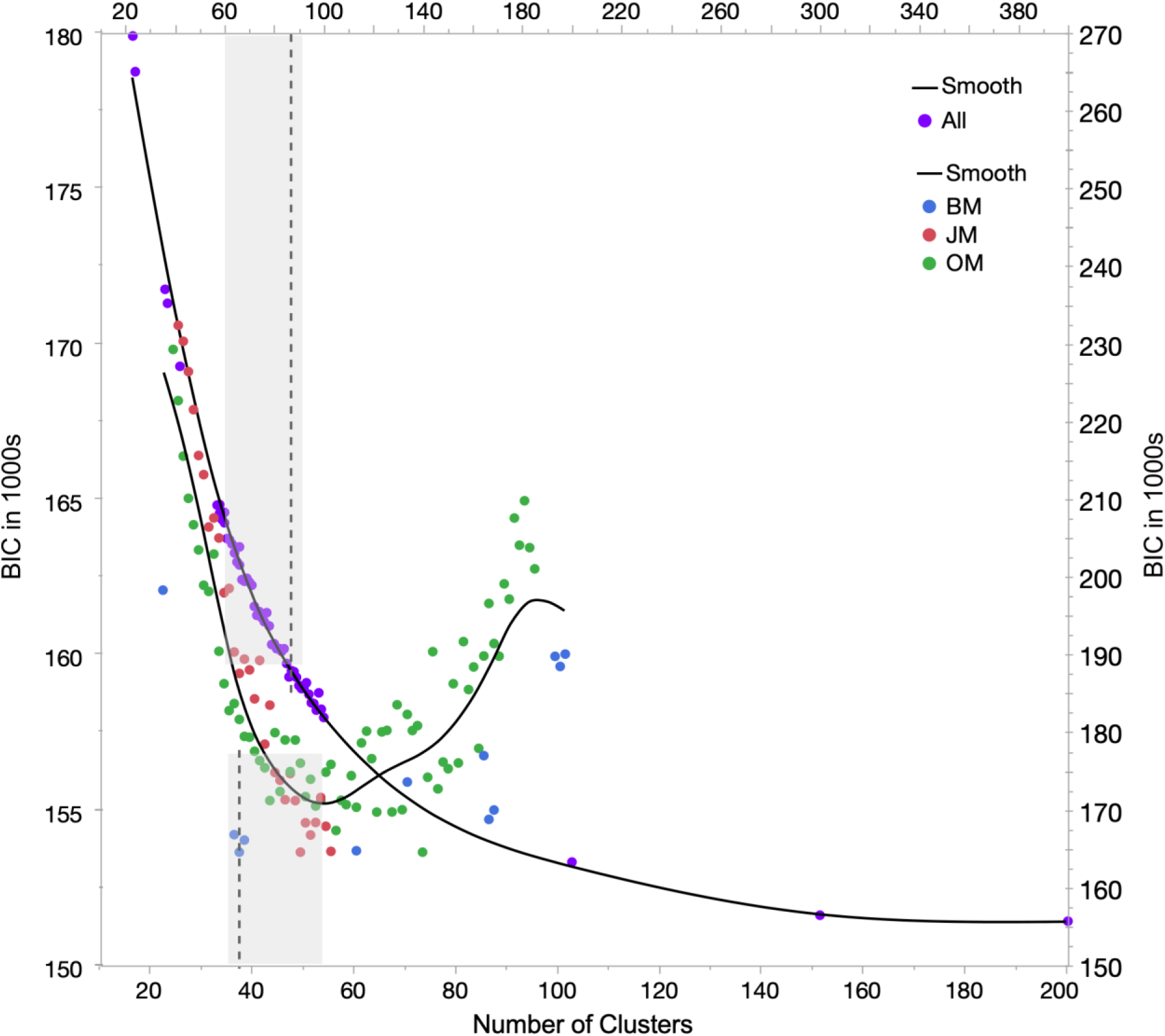
Scatterplot with a spline fit for normal mixtures cluster fit for 24 to 400 clusters for all (n = 12; purple dots) adult zebrafish sampled during 10 minutes of swimming behavior. Axes on the right shows values for BIC, and cluster numbers for this plot are labeled on the top. The left and bottom axes are used to show the range of clusters for the scatterplot with a spline fit of BIC values for a normal mixtures cluster fit obtained for three individuals (blue, green and red dots). Vertical dashed lines, show best fit for 36 clusters in a single individual (BM) and for 86 clusters for postural data pooled across all 12 animals.

### Postural clusters in a single animal

Unlike land vertebrates possessing limbs, the multivariate distribution of correlated movements of body segments in fish is relatively simple and on a lower dimensional continuum along the axial musculature. As a critical test of pose classification and to minimize the fuzziness between cluster boundaries resulting from inter-individual variation in body shape and size, we focused on the movements and postural configurations exhibited by a single animal (“BM”) that exhibited a relatively high level of swimming activity. As before, we performed a robust mixtures cluster fit using PCA seeded for 20 to 60 clusters after removing outliers. Cluster values representing the distribution of body-shapes or dynamic poses plotted as a scatterplot for the first two principal components (PCs), however, failed to exhibit well-defined boundaries (Figure 4A). Nevertheless, the normal mixtures clustering algorithm extracted 41 common postures (excluding an outlier cluster) or default body shapes that the fish reverted to as it transitioned from one action pattern to another.

**Figure 4.**
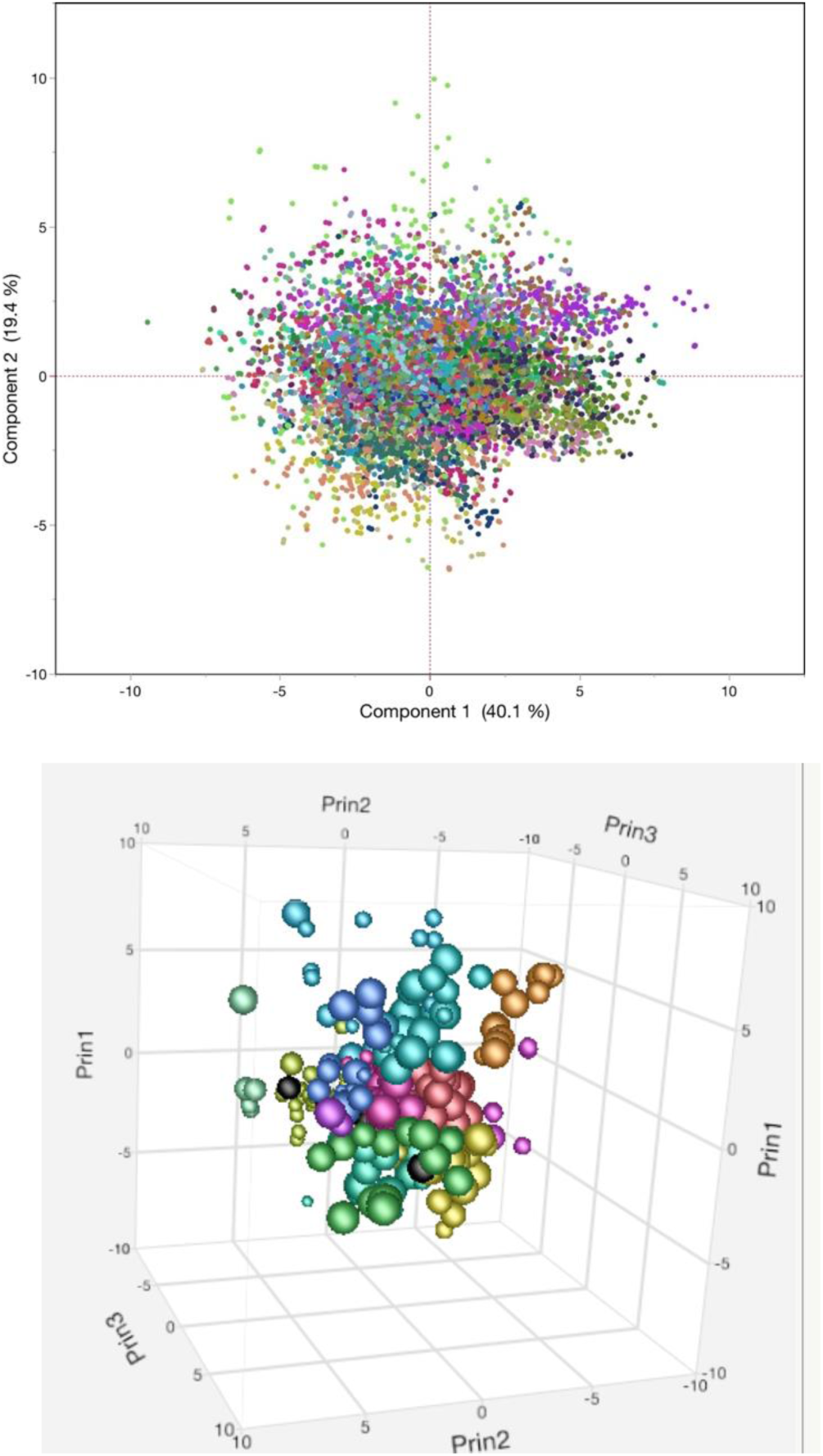
Scatterplot showing clustering of first three principal components (PCs) for the postural feature space obtained from DLC markerless tracking of an adult zebrafish underlying the different poses. Scatterplot showing clustering of first three PCs for the postural feature space after the centroids of normal mixture clusters were used to re-cluster the data. This yielded a more tight and discrete grouping of clusters within a 3D plot. Each color in the plot corresponds to a different pose.

To obtain a discrete representation of the postural configurations, we therefore used the pose feature space corresponding to the centroids of normal mixture clusters to extract the PCs underlying the different poses. These PC parameters were used to re-cluster the data (Supplementary figure 2). For fourteen of 22 uncorrelated computed parameters the normal mixtures clustering on correlations identified a best-fit (lowest Bayesian Information Criterion (BIC) value for 36 clusters for a seemingly continuous spread of configurations in multidimensional space (see Figure 3). Cluster grouping within a 3D plot of the first three PCs was dense and discreet (smallest BIC = 216634; P<0.001) (Figure 4B). Each color in the plot corresponds to a different pose. Altogether, the first three PC’s captured 80% of the variation and the first two PCs captured 70% of the variation in the pose configurations.

To visualize postural configurations, we localized the centroids to individual frames within a video sequence of body movements. Since the grouped poses represent the centroids of variants emerging from bodily motion, we label these as “dynamic poses”. The centroid of each cluster corresponded to a typical body posture along with body configurations leading into and away from that posture. A frame-by-frame analysis of the video-recording showed that the centroid of a particular dynamic pose consistently corresponded to the frames at the center of multiple examples of an action sequence or swim pattern. This indicated that the originally obtained diffuse clusters included frames grouped together because of their temporal proximity to a few central or target frames. The clusters represented a continuum of small variations overlapping with transitions either towards or away from the target pose. This realization helped us to identify and discretely classify characteristic body shapes and poses that corresponded to each cluster.

### Classification of postural configurations

To visualize the relative proximity of the 36 clusters, we performed multidimensional scaling (MDS) for the first two dimensions. MDS of the mean values for each multivariate cluster revealed a roughly circular configuration (Figure 5). The two-dimensional layout helped to visualize the relative proximity of each cluster mean to others. This analysis placed similar postures, with small variations in potentially continuous multivariate space closer to each other. On the basis of careful visual inspection, we created broader categories or superclusters of one to five closely placed cluster centers (mean values) to further condense the clustering to major inflection zones along the body axis. We discovered that each supercluster represented bends at common segments along the body axis. This supervised, manual clustering revealed an interesting design for angular flexions of the body.

**Figure 5.**
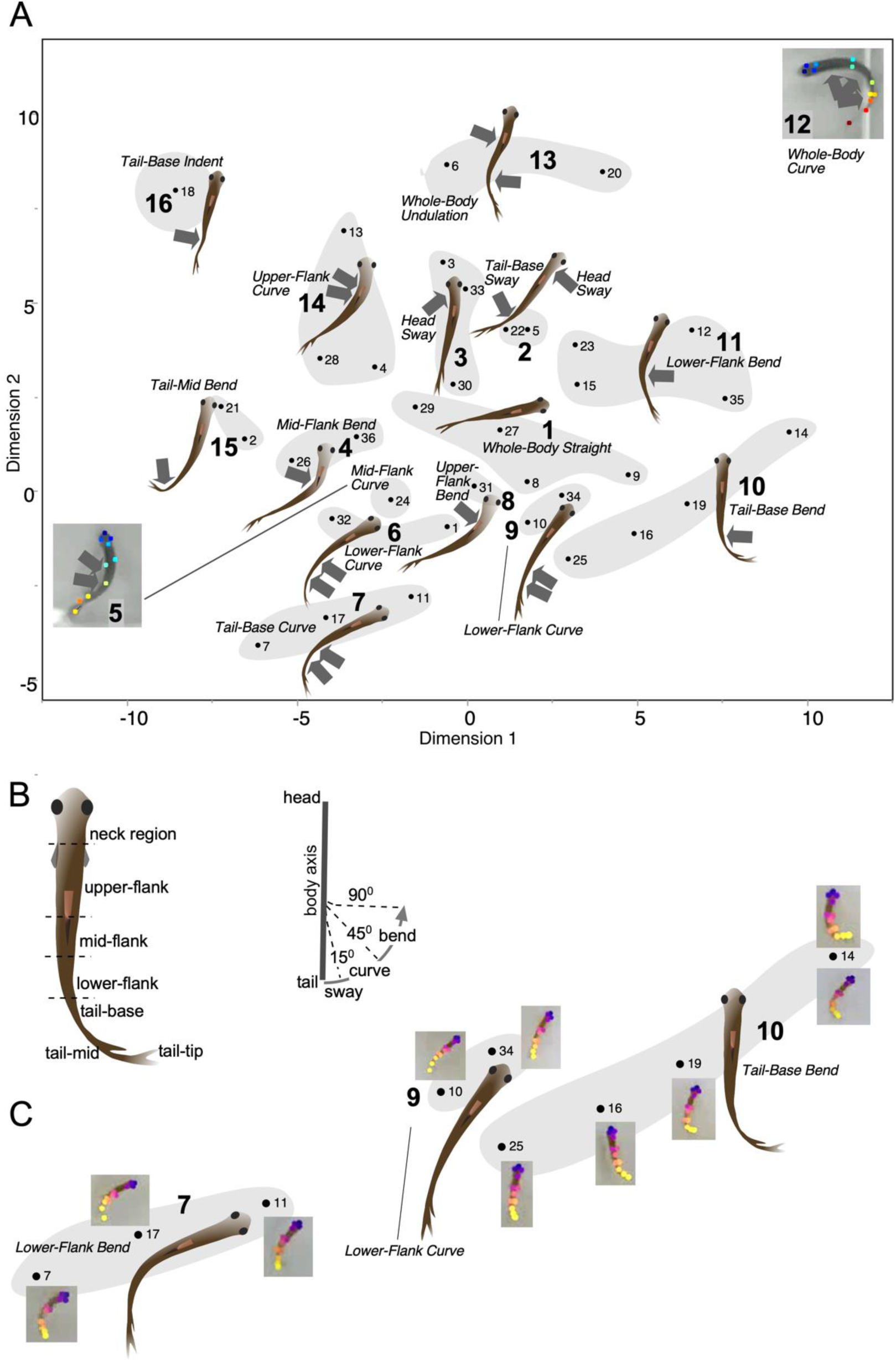
A. A multidimensional-scaled plot of the clusters of behavioral postures in a single animal extracted using a normal mixtures algorithm (JMP v.15.2, SAS Institute Inc). A best fit was found for 36 clusters that were further grouped into 12 superclusters based upon visual inspection of the general orientation of the body and upon the dominant body part involved (arrows) and its subregion as well as the general pattern/overall body shape assumed by the animal (see table 1). B. Left: dorsal view of the zebrafish body showing the three anteroposterior levels of the flank. Neck region is the anterior end of the upper-flank that includes the pectoral fins upto the anterior attachment of the pelvic fin, mid-flank covers the extent of the pelvic fin and lower-flank covers the extent of the anal fin. B:Right: diagram showing the flexion, angular flexion, defining the terms, “sway” for a gentle angular displacement of < 10°, “curve” for an angular displacement of 15° to 45° that is gradual over a relatively long flank segment, and “bend” for an angular displacement of 45° to 90°. C. Variations in dynamic poses identified by the clustering algorithm. The variations show the relative degrees to which a particular body segment could be flexed, leading to a variety of poses associated with different action patterns as well as special maneuvers exercised during swimming.

**Table 1.**
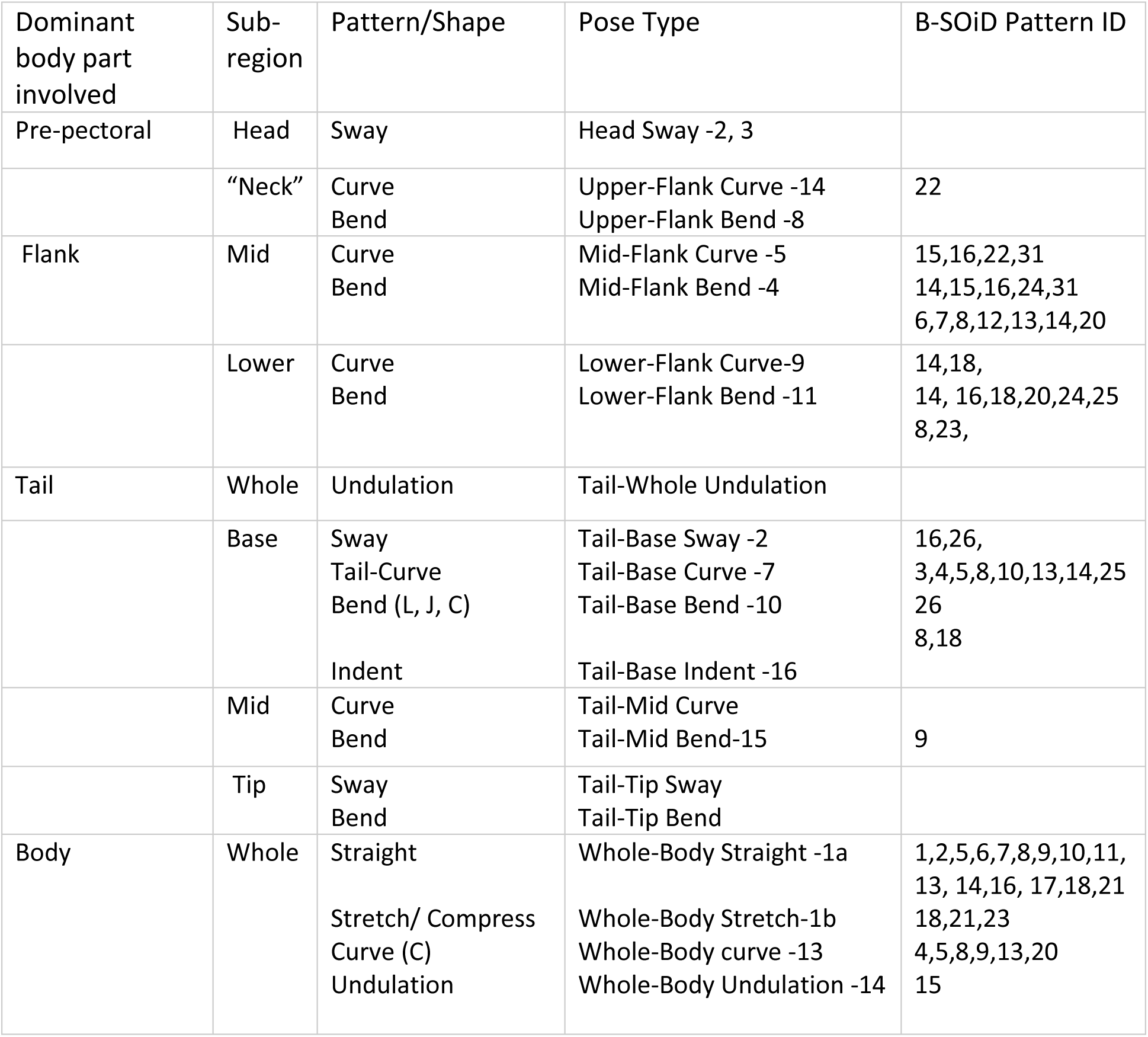
Body parts and postural body congiurations used for pose classification in zebrafish. Body parts and subregions involved in movement were considered together with the geometric pattern in developing a terminology to classify different poses. The tip of the tail was not included as it was frequently invisible and underwent rapid, large and generally rhythmic displacements during free-swimming and turning.

| Dominant body part involved | Sub-region | Pattern/Shape | Pose Type | B-SOiD Pattern ID |
| --- | --- | --- | --- | --- |
| Pre-pectoral | Head | Sway | Head Sway -2, 3 |  |
|  | “Neck” | Curve<br>Bend | Upper-Flank Curve -14<br>Upper-Flank Bend -8 | 22 |
| Flank | Mid | Curve<br>Bend | Mid-Flank Curve -5<br>Mid-Flank Bend -4 | 15,16,22,31<br>14,15,16,24,31<br>6,7,8,12,13,14,20 |
|  | Lower | Curve<br>Bend | Lower-Flank Curve-9<br>Lower-Flank Bend -11 | 14,18,<br>14, 16,18,20,24,25<br>8,23, |
| Tail | Whole | Undulation | Tail-Whole Undulation |  |
|  | Base | Sway<br>Tail-Curve<br>Bend (L, J, C)<br>Indent | Tail-Base Sway -2<br>Tail-Base Curve -7<br>Tail-Base Bend -10<br>Tail-Base Indent -16 | 16,26,<br>3,4,5,8,10,13,14,25<br>26<br>8,18 |
|  | Mid | Curve<br>Bend | Tail-Mid Curve<br>Tail-Mid Bend-15 | 9 |
|  | Tip | Sway<br>Bend | Tail-Tip Sway<br>Tail-Tip Bend |  |
| Body | Whole | Straight<br>Stretch/ Compress<br>Curve (C)<br>Undulation | Whole-Body Straight -1a<br>Whole-Body Stretch-1b<br>Whole-Body curve -13<br>Whole-Body Undulation -14 | 1,2,5,6,7,8,9,10,11,<br>13, 14,16, 17,18,21<br>18,21,23<br>4,5,8,9,13,20<br>15 |

For a systematic description of the condensed postural design, we developed a nomenclature for different poses by naming bends at the main antero-posterior level of the body axis and describing the pattern or shape achieved given the degree of bending observed. This idealized classification scheme used consistently for bends at all body segments resulted in 18 possible dynamic poses that we believe capture the major postural configurations possible in adult zebrafish (see table 1). Three of the possible poses per this scheme were not observed in the single animal data. Most poses could be defined by a bend at middle and lower segments of the flank. Some poses, such as upper and lower body curves, consisted of a slight bend at a combination of locations, yielding a smooth flexion of the body. At the rostral end, only a slight bend of the rostral-most flank portion yielded a “Head-sway”. The base of the tail was a particularly flexible locus. The “straight body” pose was typically associated either with a stationary position or a longitudinal stretching or compression of the body. The characteristic C-bend of the startle reflex is not included given its rare occurrence though it is similar to the whole-body curve (Supercluster # 12 in Figure 5). Note that the analysis involving the entire cohort of animals tested yielded a larger number of clusters but pose configurations were not extracted manually in each case as this is a relatively laborious process.

### However, B-SOiD-extracted configurations revealed the presence of similar poses

As a further test of the distinctness of each supercluster, we pooled the mean parametric values of each of the statistically identified clusters contained within superclusters, and performed a principal component analysis that revealed a clear segregation of each cluster type. There may be additional categorically different poses not observed here, e.g., fish rotation of the body by ∼ 90° (Kanwal: personal observation), but we believe our analysis captured the majority of the commonly observed poses within a two-dimensional horizontal plane of swimming. We did not perform a detailed analysis of potential poses exhibited when fish swim up and down the depth dimensional of the water. However, from casual observations, these poses mostly represent the elevational body angle and rate of tail fin undulations corresponding to the speed at which fish ascend and descend in the water. It is not expected to produce new body shapes that are not observed in the two-dimensional horizontal plane.

Pose analysis unexpectedly also revealed compression of ∼8% of body length (BL) and stretch of ∼4% of BL, as in stretch/compression of an earthworm’s body along its longitudinal axis. Whole body compressions could play an important role in reaching actions and in streamlining the body to minimize lateral forces on the flank during turning movements or when passing through a narrow opening.

## Discussion

Fish behavior has been studied previously from ecological, social and evolutionary perspectives. Zebrafish display both interesting behavior and easily accessible neural processes [Levin, and Cerutti, 2009]. Here we focused on postural patterns to gain a better understanding of the evolutionary design of movement and its control in vertebrates. An efficient movement strategy is of fundamental importance to the success of all vertebrates in water, on land and in the air [Burgess, and Granato, 2008]. Rapid advances in imaging techniques and most recently in ML-powered behavior-tracking are facilitating an understanding of behavior from a structural-functional perspective and eventually as an output of specific neural circuits and genes [Kane et al., 2020; Mathis et al., 2018]. As a first step towards advancing this understanding, we classified consistently identifiable postures exhibited by free-swimming adult zebrafish. Despite the continuous and relatively smooth transitions of body configurations, we were able to identify key postures that corresponded to statistically significant occurrences of relatively stable inflection loci during swimming and turning maneuvers. We postulate that these small number of identifiable body configurations or “dynamic poses” represent target postures which constitute key elements for the organization of behavior. They may represent functional units of behavior for elaborating temporally complex constructs. By this scheme, neural circuits are designed to contract muscle sets to achieve well-defined target postures, and behavior is a natural outcome of the mechanics resulting from reaching a target posture or enacting a sequence of target poses. In short, the inflection points and associated bodily configurations, though momentary, are consistently observed patterns, constituting the center of a cluster of contiguous pose variants approaching towards and receding from a target pose. Identifying these target poses may constitute a key step in elucidating the neural circuits and/or multi-neuronal activity patterns associated with achieving a particular pose.

### Classification of swimming dynamics and movement postures

An earlier study attempted to classify short term movement patterns in adult zebrafish [Liu et al., 2011]. Although a method using self-organizing maps and hidden Markov processes was developed and applied, no clear classification other than short-term stop-and-go patterns, zigzag movements, turns and backward swimming was noted [Liu et al., 2011; Li et al., 2013]. That methodology was not designed to reveal postures; longer-term behaviors were expected to be identified in a later study, which was apparently not conducted. From our examination, the locations of bending in the most common postural configurations in adult zebrafish were largely constrained to three locations – the upper, mid and lower flank. Although axon terminations of reticulospinal neurons on motor neurons are present all along the spinal cord, constraining the activation and muscle contraction to one axial level on either side can pattern postural configurations, such as J-bend, C-bend, etc., that help with navigation around objects while swimming or simply to make different types of turns, including a U-shape to turn around. Localizing of body bends to particular regions of the body was intriguing because axial musculature and motor neuron innervation is present throughout the body. A close examination of serial sections at different levels of the spinal cord may identify differences in synaptic densities or relative sizes of muscle groups to establish a neuromuscular basis for localization of movement control.

The dynamics of body postures in fish involve stabilization and maximization of the efficiency of undulating movements of the body with respect to muscle contraction forces acting on the vertebrae (body stiffness) and fluid dynamics, such as buoyancy, water resistance and flow, for both turning and swimming [Torigoe et al., 2021; Tytell et al., 2014]. Bending of the flank is a complex interaction of bone, cartilage, a segmented muscle design, and ligaments, all working together. Lateral flexion in fishes is quite different from the lateral flexion seen in mammals and is essential for various activities, including swimming, maneuvering, escaping predators, and hunting prey. Experiments demonstrate that largemouth bass possess the capability to increase body stiffness by using their muscles to generate negative mechanical work during an oscillatory pattern of bending of a ribbon-like body [Long, and Nipper, 1996; Lauder et al., 2011]. Therefore, the target poses associated with partial undulation may result from changes in the stiffness properties of the flank with greater contraction at one or more specific loci along the vertebral column. One example of an important type of movement exhibited by both larvae and adults is the startle response. A similar mechanism exists in most vertebrate species, including humans where it manifests itself in a number of ways, including blinking and involuntary limb bending (Poli & Angrilli, 2015). In fish, startle responses have been categorized into two classes, a C-start that is exhibited by both larvae and adults, and a S-start that only occurs in larvae. A C-start response involves the sharp curving of the body of the fish (or larva) towards one side in the shape of a “C”, as the name suggests. This response is mediated by Mauthner cells [Eaton, 1977].

Swimming movements in adult zebrafish are continuous, requiring a constant output of energy whereas larvae swim in a “burst and glide” manner, using a short, explosive initial burst of energy to propel them to glide afterwards followed by the next burst [Berg et al., 2018; Johnson et al., 2020]. Previous studies of pose classification in zebrafish larvae, captured multi-timescale structure from behavioral data by developing computational tools to track continuously across multiple 24-h day/night cycles. They extracted millions of samples of movements and pauses, termed bouts, and used unsupervised learning to reduce each larva’s behavior to an alternating sequence of active and inactive bout types, termed modules [Ghosh, and Rihel, 2020]. Through hierarchical compression, they also identified recurrent behavioral patterns or motifs. They reported that the duration of both modules and motifs, as defined, varied across the day/night cycle, revealing behavioral structure at sub-second to day-long timescales. Their analysis, however, did not capture data to allow a classification at the level of dynamic poses.

### Body-stretch and compression

Some of the universal patterns of translational movement in animals involve either compression and stretch, as commonly observed in earthworms, undulatory or sinusoidal movements as typically observed in the aquatic environment, and anchored movement involving a fulcrum, from which the rest of the body can be leveraged to move either forward or backward or sideways [Ozkan-Aydin et al., 2021; Bicanski et al., 2013; Cabelguen et al., 2003]. Both invertebrate and vertebrate species use limbs to move in this way. An interesting observation from close examination of markerless tracking was that zebrafish and possibly other fish species can extend and compress their body in addition to the transverse bends. Flank musculature in fish species is anatomically organized as W-shaped bands that were first described several decades ago [Romer, 1977]. This angular orientation of the flank musculature offers an interesting design feature where the same muscles can be used both for longitudinal contractions of the body as well as being effective for transverse bends. In other words, they present an optimization of the longitudinal and circular muscle design present in the gut undergoing peristaltic movements, and in the body wall of earthworms, allowing them to move along substrate by alternate contraction and relaxation of longitudinal and circular muscles together with setae that can grasp onto substrate roughness [Mizutani et al., 2004; Tanaka et al., 2012; Ozkan-Aydin et al., 2021].

### Neural control of posture-driven action

Quantitative studies of behavior can provide a noninvasive window into its neural control and eventually its genetic underpinnings [Sumbre, and de Polavieja, 2014]. Initiation of turning and swimming movements may originate with neural activity in the brainstem or at higher levels of the brain and get translated into a set of actions controlled by the lower motor system [Benhamou, 2010]. Temporal patterns of neural activity contract and relax antagonistic and other sets of muscles synchronously and/or sequentially [Saint-Amant, and Drapeau, 1998]. Though exterosensory inputs can create variations in the expression of locomotory movements, the default output pattern for swimming maneuvers is thought to be programmed within a special type of neural network known as a motor pattern generator or more generically as a central pattern generator (CPG) [Marder, and Bucher, 2001; Stein, 2005]. Motor pattern-related CPGs are generally present within the brainstem and spinal cord of vertebrates [Sweeney, and Kelley, 2014; Buchanan, 2018; Travers et al., 1997]. These CPGs produce rhythmic patterns of contractions in relevant muscle groups. Together with proprioceptive feedback [Hamlet et al., 2018], they determine the timing, phasing, and intensity parameters to drive motoneuron output in the absence of higher order descending control that can transiently bias the output of CPGs [Severi et al., 2014]. We briefly discuss here how our results reflect back on the neural and mechanical aspects of postural control.

Mechanisms contributing to posture and trajectory control can be either passive or active [Gordon et al., 2008]. Passive mechanisms operate automatically and continuous whereas active i.e., those powered by neural activity, drive movements of body and fin (or limb in land vertebrates). They require active coordination and control. Coordination of changes in body stiffness, resulting in behaviorally and dynamically relevant body postures, must require intricate neural signaling between body stiffness- and curvature-based proprioceptive feedback, vestibular activity and CPGs [Hamlet et al., 2018]. Establishing the geometry of dynamic poses allows us to think of continuous motion and smooth transitions between different behaviors as being driven by multiple CPGs that interact to facilitate a switch between different target poses. For example, a transition between Lower-Flank Curve and Lower-Flank Bend or between Whole-Body Undulation and Upper-Flank Curve poses (see figure 5A). Poses achieved during directional turns in zebrafish may be functionally equated to reaching arm movements in 3D space in primates via a population of directionally tuned neurons [Georgopoulos et al., 1988]. Moreover, the same target in 3D space can be achieved in multiple ways by differentially activating central neural circuits, such as in the basal ganglia and cortex, and relevant muscle groups to achieve different arm trajectories [Lee, and Assad, 2003]. In zebrafish, the transition between two target poses or their termination may depend upon a specific form of body curvature-based, closed-loop proprioceptive feedback that signals the achievement of a particular pose as in lampreys [Hamlet et al., 2018]. The target poses themselves may be encoded in the activity of higher-order single neurons or local circuits within the motor cortex and basal ganglia [Hermer-Vazquez et al., 2004] that pre-shape postures and drive lower motor neurons through the CPGs. How exactly CPG activity contracts flank musculature at specific locations while the rest of the body simply bends via a stiffness gradient along the rest of the flank musculature in adult zebrafish needs further examination of electromyographic and neural activity patterns along the spinal cord. The specific genes associated with the construction of CPGs during development have not been clearly identified or manipulated [Marder, and Rehm, 2005], but these can be easily studied in zebrafish by administering drugs blocking gene expression directly through the aquatic medium.

### From water to land: a perspective on the evolutionary advancement of postural design

Movement patterns associated with locomotion are particularly important for exploration, navigation, and social interactions in all animal species, and can involve complex sequences of body and limb trajectories. We have shown that various forms of turning movements are characterized by postural configurations resulting from bends at mainly three anteroposterior levels. The neck region, at the anterior border of the upper flank in adult zebrafish, is accentuated as a slender, flexible segment in land animals. It facilitates head orientation for maintaining gaze and for initiation of directional turns. The mid- and caudal-flank locations correspond to the levels that generate stepping postures in land vertebrates. The base of the tail-fin at the caudal end of the vertebral column is a fourth locus of bending that corresponds to the base of the tail in land animals. In zebrafish, the base of the tail acts as a fulcrum for undulating tailfin movements that can propel the fish forward at extremely high speeds. (Plaut, 2000). Tail movements in tetrapods allow for stability during locomotion over ground and when performing arial acrobatics and during gliding in arboreal species.

From a biomechanics perspective, the fish-like design of bending at mid and caudal locations along the body axis was well suited for turning and slithering type of movements on a wet substrate as fish-like creatures evolved to adapt to land (Figure 6). As evidence, the mechanics of mode-switching in limbless and legged animals follow a common pattern [Kuroda et al., 2014]. The same holds for swimming in aquatic species. During overground stepping in amphibious animals, such as adult newts, there is a correspondence between rhythmic activation of muscles within the limb and those along the body axis for swimming [Katz 2016]. The pattern of rhythmic activation of epaxial myomeres during stepping and swimming in amphibious species also match a hybrid form of their activation for swimming in lampreys [Delvolvé et al., 1997]. In short, limbless crawlers first moved in a manner similar to walking before the evolution of legs [Kuroda et al., 2014]. Thus, the existence of only two pairs of limbs in land vertebrates, as compared to six in insects and more in other invertebrate species, may be a direct outcome of the neural and biomechanical design for movement in the aquatic environment. In fact, some fish species, like *Polypterus*, were already walking under water before the evolution of tetrapods [King et al., 2011; Schuchmann, and Siemers, 2010; Hale et al., 2022].

**Figure 6A.**
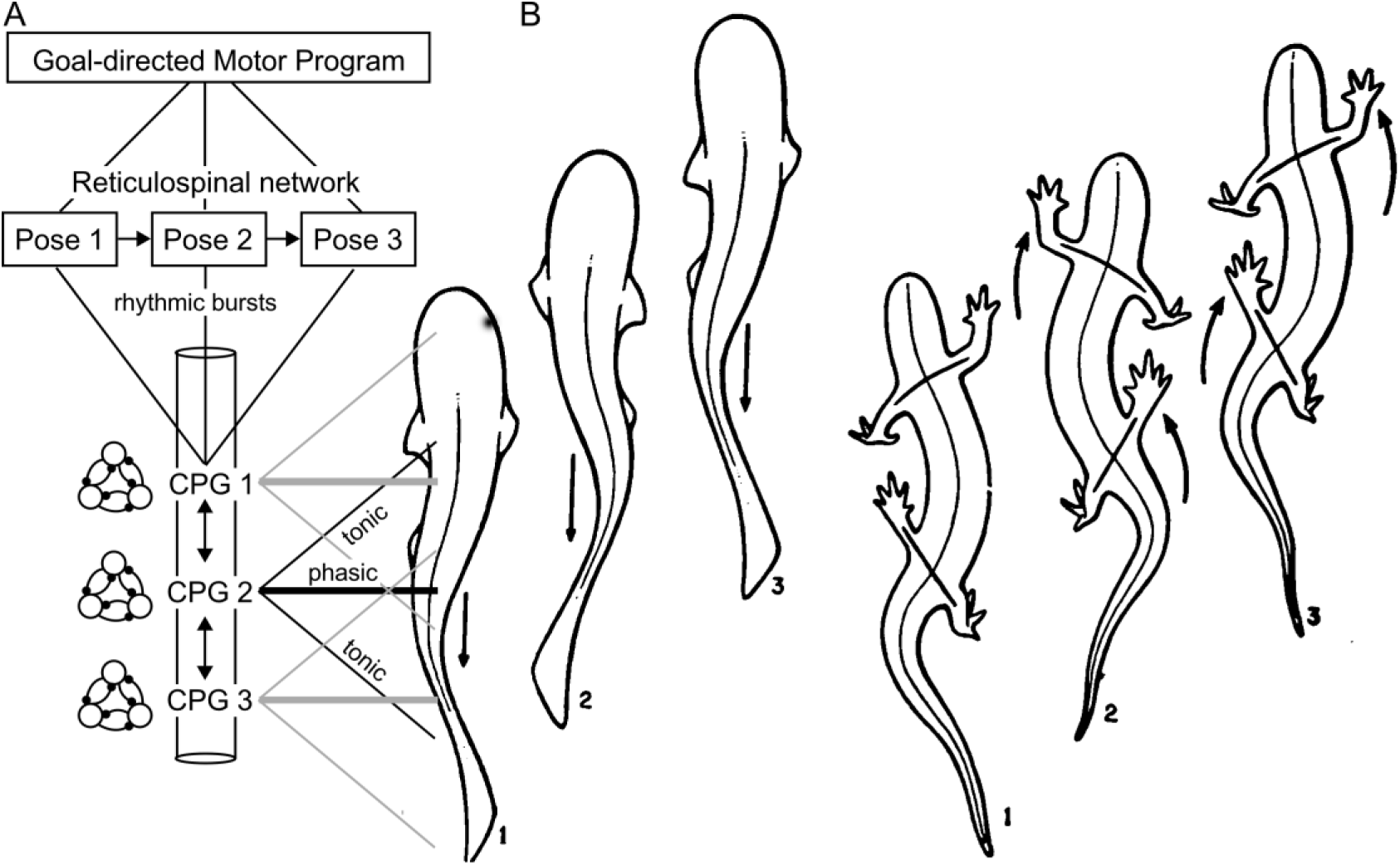
Schematic of reticulospinal networks generating differential activity within CPGs in the spinal cord to iniate pose-driven movement patterns in adult zebrafish. Experimental data support the presence of rhythmic discharges within adjacent CPGs. Phasic firing of CPG neurons (bold lines) iniates bending via contraction of myomeres present within a restricted segment of the flank. Simultaneous tonic activity (thin lines) in spinal motor output initiates stiffness changes at other levels of the flank musculature to facilitate bending at specific loci. In zebrafish, there are two classes of motor neurons (MNs) involved in CPG-controlled swimming, primary MNs and secondary MNs [Babin et al., 2014]. For postural control in adult zebrafish, the primary MN component of CPGs may trigger muscle contractions at specific loci for driving target postures whereas secondary MNs may mediate relatively nonspecific changes in muscle tone leading to changes in body stiffness. All of this activity is also under the control of vestibular and proprioceptive feedback, e.g., from muscle spindles controlling muscle tension, facilitating intersegmental coordination by phase coupling [Ijspeert, 2001; McPherson et al., 1994; Delvolvé et al., 1997; Grillner et al., 2017; Cohen, 1987]. B. Dorsal view of body flexions associated with locomotion in a fish (a) vs. a salamander (b) (Romero, 1977). As illustrated here, the primary nodes of bending in fish and the control of limbs at their proximal end correspond to discrete upper and lower regions on the flank.

In summary, an analysis of zebrafish movement indicated its organization as a set of twelve identifiable “dynamic pose” categories that included stretch and compression of the body. Pose configurations revealed discrete flexion zones along the anteroposterior levels of the flank that predominantly account for the generation of different poses. Target postures were constrained to bends at two anteroposterior levels of the body axis with a third level near the head exhibiting minor bends. Description and classification of the transitional patterns connecting target postures is the focus of a follow-up study (Sanghera and Kanwal, in preparation). We argue that the musculoskeletal design for bending at predominantly two mid-to-posterior loci together with passive undulatory dynamics for body flexion accounts for the emergence of fore- and hind-limbs for laterally directed turns and locomotion (Romer, 1977). The emergence of digits or claws at the distal end of these limbs allowed animals to grasp the rough substrate for translating the body forward as well to shift the direction of movement. Comparative studies of musculoskeletal analysis in conjunction with neurophysiological recordings from spinal cord CPGs are needed to test these insights resulting from our study and the work of many others.

## Acknowledgements

Supported in part by Grant #PAR3152016-003, Pioneer Academics LLC to J.S.K. from Pioneer Academics, LLC, a grant from Academic Enhancement Research Fellowship, University of Miami, to B. S. and Georgetown Zebrafish Shared Resource, supported in part by NIH/NCI grant P30-CA051008.

## Supplementary Table and figures

**Table S1:**
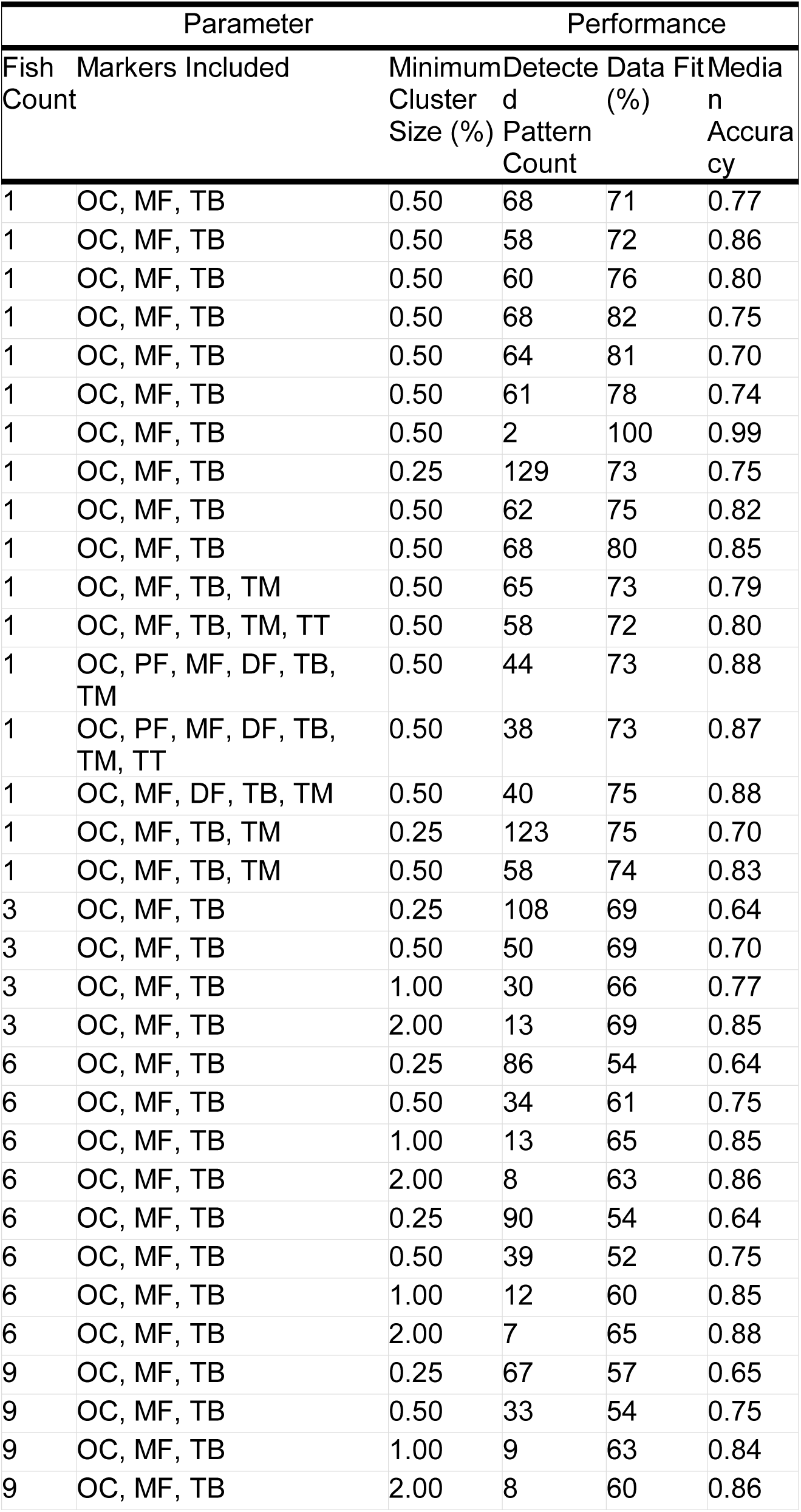
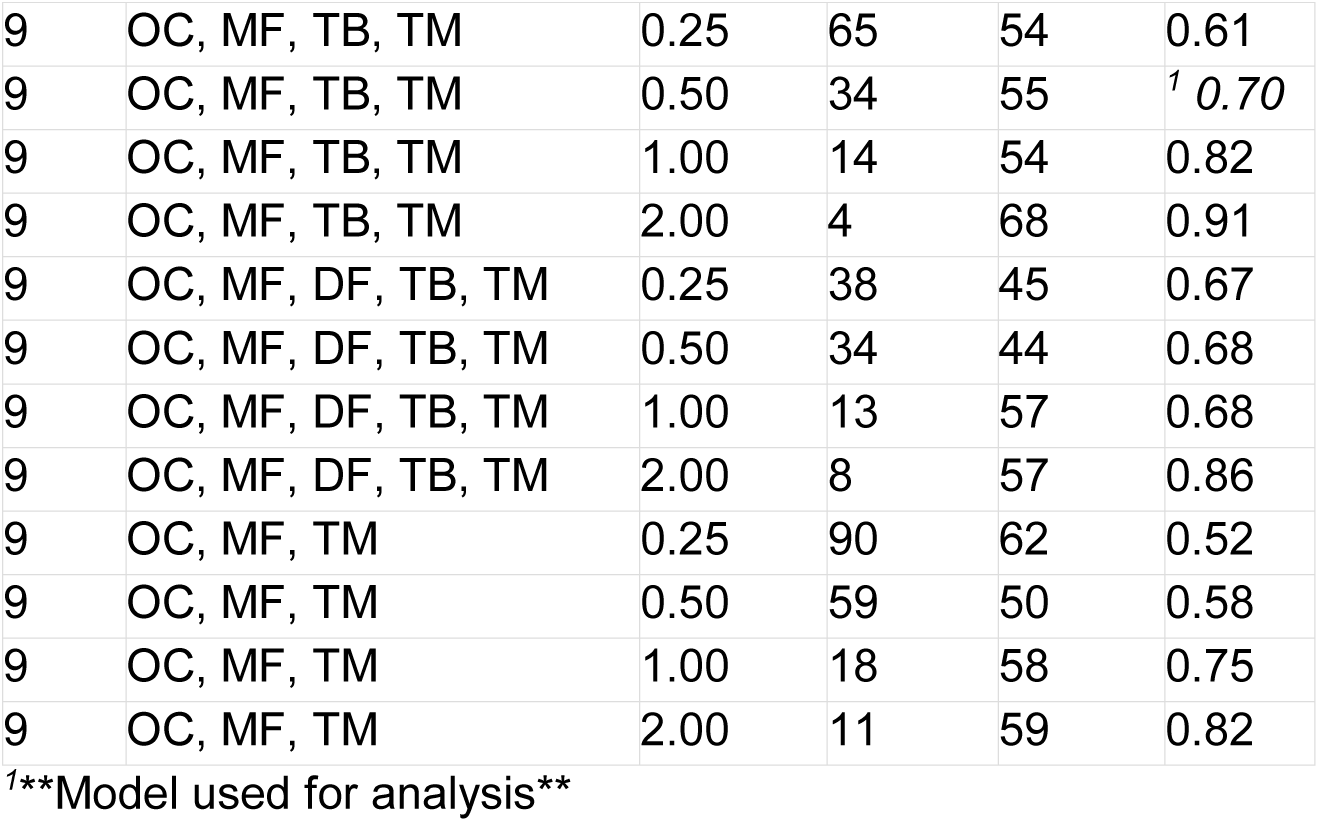
B-SOiD Models Tested as Generated from different combinations of fish, cluster size, and axis markers. Different combinations of B-SOiD parameters were used to generate the unsupervised classifier. This table lists the different combinations of fish, tracking markers, and cluster sizes used to generate the used model. The markers included dorsal fin (DF), Mid-Flank (MF), Pectoral Fin (PF), Ocular Center (OC), Tail-Base (TB) and Tail-Mid (TM).

**Supp. Figure 1.**
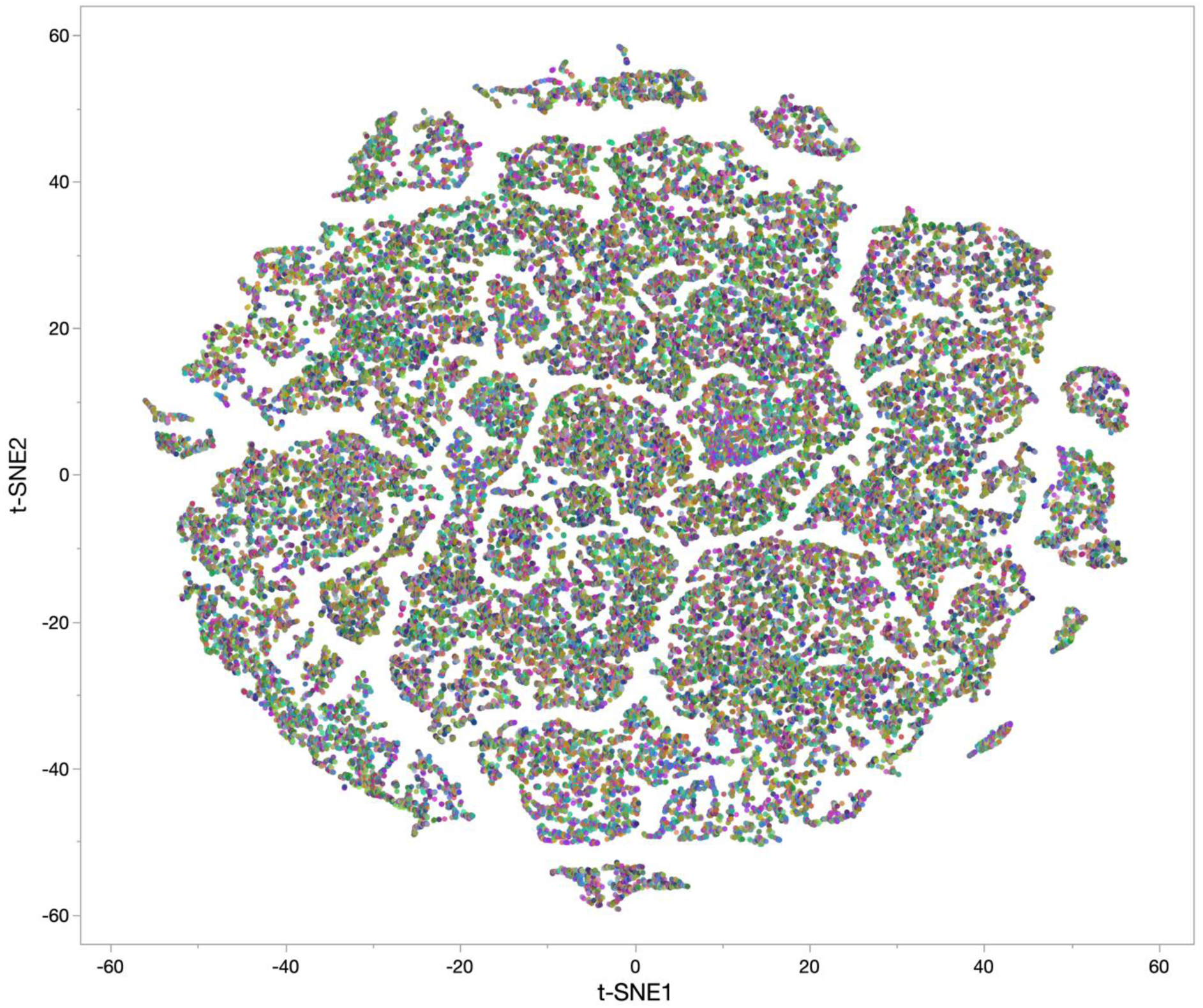
A t-SNE plot of the feature space across all animals (n = 12) with B-SOiD generated pattern ID as the color code.

**Supp. Figure 2.**
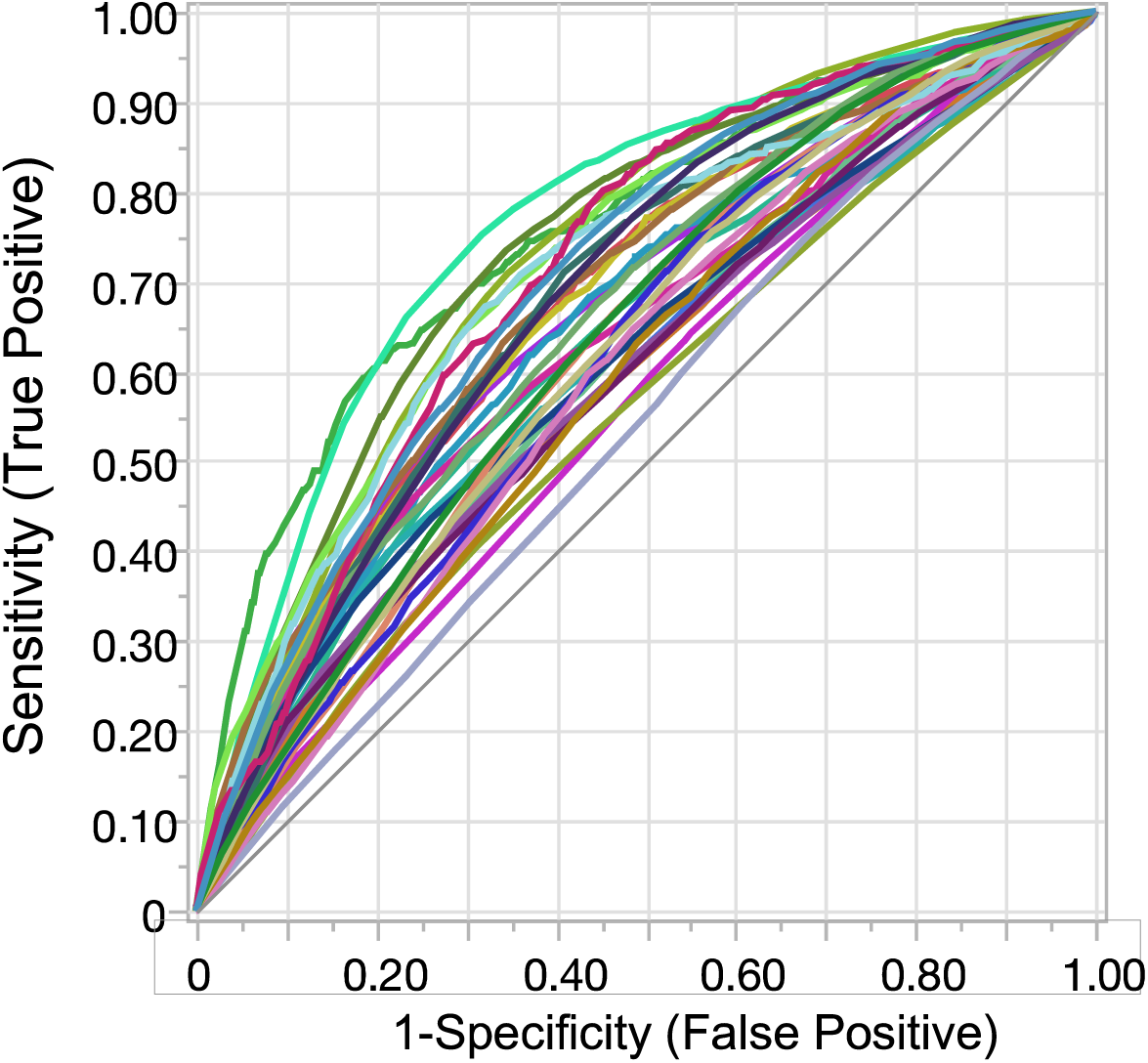
ROC curves, to show the reliability of classification based on discriminant function analysis over the first three minutes of swimming with pattern ID as the classifier (shown as different colors). Proximity of the curves to the diagonal indicates a lower reliability of fit (smaller area under the curve) to a particular ID.

