## Supplemental Table 1 for "Pose analysis in free-swimming adult zebrafish, *Danio rerio*: “fishy” origins of movement design"

### Supplementary Table and figures

| Parameter | | | Performance | | |
| --- | --- | --- | --- | --- | --- |
| Fish Count | Markers Included | Minimum Cluster Size (%) | Detected Pattern Count | Data Fit (%) | Median Accuracy |
| 1 | OC, MF, TB | 0.50 | 68 | 71 | 0.77 |
| 1 | OC, MF, TB | 0.50 | 58 | 72 | 0.86 |
| 1 | OC, MF, TB | 0.50 | 60 | 76 | 0.80 |
| 1 | OC, MF, TB | 0.50 | 68 | 82 | 0.75 |
| 1 | OC, MF, TB | 0.50 | 64 | 81 | 0.70 |
| 1 | OC, MF, TB | 0.50 | 61 | 78 | 0.74 |
| 1 | OC, MF, TB | 0.50 | 2 | 100 | 0.99 |
| 1 | OC, MF, TB | 0.25 | 129 | 73 | 0.75 |
| 1 | OC, MF, TB | 0.50 | 62 | 75 | 0.82 |
| 1 | OC, MF, TB | 0.50 | 68 | 80 | 0.85 |
| 1 | OC, MF, TB, TM | 0.50 | 65 | 73 | 0.79 |
| 1 | OC, MF, TB, TM, TT | 0.50 | 58 | 72 | 0.80 |
| 1 | OC, PF, MF, DF, TB, TM | 0.50 | 44 | 73 | 0.88 |
| 1 | OC, PF, MF, DF, TB, TM, TT | 0.50 | 38 | 73 | 0.87 |
| 1 | OC, MF, DF, TB, TM | 0.50 | 40 | 75 | 0.88 |
| 1 | OC, MF, TB, TM | 0.25 | 123 | 75 | 0.70 |
| 1 | OC, MF, TB, TM | 0.50 | 58 | 74 | 0.83 |
| 3 | OC, MF, TB | 0.25 | 108 | 69 | 0.64 |
| 3 | OC, MF, TB | 0.50 | 50 | 69 | 0.70 |
| 3 | OC, MF, TB | 1.00 | 30 | 66 | 0.77 |
| 3 | OC, MF, TB | 2.00 | 13 | 69 | 0.85 |
| 6 | OC, MF, TB | 0.25 | 86 | 54 | 0.64 |
| 6 | OC, MF, TB | 0.50 | 34 | 61 | 0.75 |
| 6 | OC, MF, TB | 1.00 | 13 | 65 | 0.85 |
| 6 | OC, MF, TB | 2.00 | 8 | 63 | 0.86 |
| 6 | OC, MF, TB | 0.25 | 90 | 54 | 0.64 |
| 6 | OC, MF, TB | 0.50 | 39 | 52 | 0.75 |
| 6 | OC, MF, TB | 1.00 | 12 | 60 | 0.85 |
| 6 | OC, MF, TB | 2.00 | 7 | 65 | 0.88 |
| 9 | OC, MF, TB | 0.25 | 67 | 57 | 0.65 |
| 9 | OC, MF, TB | 0.50 | 33 | 54 | 0.75 |
| 9 | OC, MF, TB | 1.00 | 9 | 63 | 0.84 |
| 9 | OC, MF, TB | 2.00 | 8 | 60 | 0.86 |
| 9 | OC, MF, TB, TM | 0.25 | 65 | 54 | 0.61 |
| 9 | OC, MF, TB, TM | 0.50 | 34 | 55 | *^1^ 0.70* |
| 9 | OC, MF, TB, TM | 1.00 | 14 | 54 | 0.82 |
| 9 | OC, MF, TB, TM | 2.00 | 4 | 68 | 0.91 |
| 9 | OC, MF, DF, TB, TM | 0.25 | 38 | 45 | 0.67 |
| 9 | OC, MF, DF, TB, TM | 0.50 | 34 | 44 | 0.68 |
| 9 | OC, MF, DF, TB, TM | 1.00 | 13 | 57 | 0.68 |
| 9 | OC, MF, DF, TB, TM | 2.00 | 8 | 57 | 0.86 |
| 9 | OC, MF, TM | 0.25 | 90 | 62 | 0.52 |
| 9 | OC, MF, TM | 0.50 | 59 | 50 | 0.58 |
| 9 | OC, MF, TM | 1.00 | 18 | 58 | 0.75 |
| 9 | OC, MF, TM | 2.00 | 11 | 59 | 0.82 |
| *^1^***Model used for analysis** | | | | | |

Table S1: B-SOiD Models Tested as Generated from different combinations of fish, cluster size, and axis markers. Different combinations of B-SOiD parameters were used to generate the unsupervised classifier. This table lists the different combinations of fish, tracking markers, and cluster sizes used to generate the used model. The markers included dorsal fin (DF), Mid-Flank (MF), Pectoral Fin (PF), Ocular Center (OC), Tail-Base (TB) and Tail-Mid (TM).
