## Supplemental Fig 1 for "Pose analysis in free-swimming adult zebrafish, *Danio rerio*: “fishy” origins of movement design"

### Supplementary figure 1

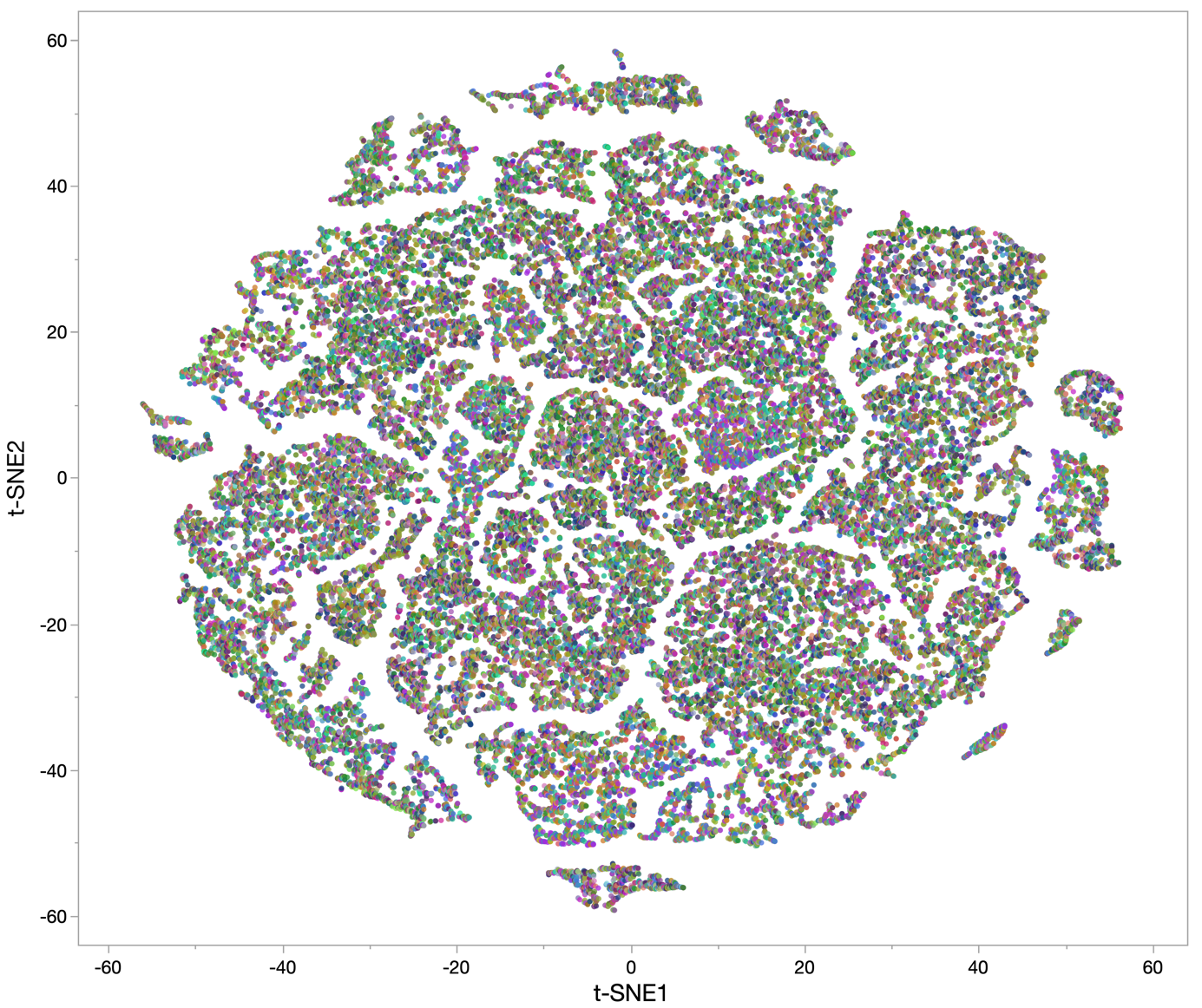

Supp. Figure 1. A t-SNE plot of the feature space across all animals (n = 12) with B-SOiD generated pattern ID as the color code.
