## Supplemental Fig 2 for "Pose analysis in free-swimming adult zebrafish, *Danio rerio*: “fishy” origins of movement design"

### Supplementary figure 2

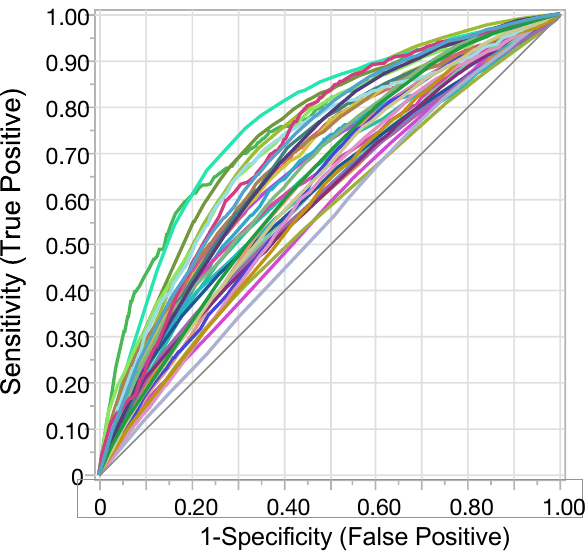

Supp. Figure 2. ROC curves, to show the reliability of classification based on discriminant function analysis over the first three minutes of swimming with pattern ID as the classifier (shown as different colors). Proximity of the curves to the diagonal indicates a lower reliability of fit (smaller area under the curve) to a particular ID.
